# A Comparison of Non-human Primate and Deep Reinforcement Learning Agent Performance in a Virtual Pursuit-Avoidance Task

**DOI:** 10.1101/567545

**Authors:** Theodore L. Willke, Seng Bum M. Yoo, Mihai Capotă, Sebastian Musslick, Benjamin Y. Hayden, Jonathan D. Cohen

## Abstract

We compare the performance of non-human primates and deep reinforcement learning agents in a virtual pursuit-avoidance task, as part of an effort to understand the role that cognitive control plays in the deeply evolved skill of chase and escape behavior. Here we train two agents, a deep Q network and an actor-critic model, on a video game in which the player must capture a prey while avoiding a predator. A previously trained rhesus macaque performed well on this task, and in a manner that obeyed basic principles of Newtonian physics. We sought to compare the principles learned by artificial agents with those followed by the animal, as determined by the ability of one to predict the other. Our findings suggest that the agents learn primarily 1st order physics of motion, while the animal exhibited abilities consistent with the 2nd order physics of motion. We identify scenarios in which the actions taken by the animal and agents were consistent as well as ones in which they differed, including some surprising strategies exhibited by the agents. Finally, we remark on how the differences between how the agents and the macaque learn the task may affect their peak performance as well as their ability to generalize to other tasks.

## 1 Introduction and background

There is mounting interest in comparing the performance of artificial agents with natural agents in complex task domains. While in some domains, artificial agents have been successful in outperforming humans (*1–3*), there are other domains in which they still fail to do so (*4*). On the one hand, the systematic comparison of artificial and natural agents can help identify weaknesses in state-of-the-art artificial intelligence (AI) and, on the other hand, it can help to better understand the cognitive processes, and underlying neural mechanisms responsible for the behavior of natural agents, including humans. However, comparisons between artificial and natural agents are often limited to crude behavioral measures, such as overall task performance, and are constrained by a lack of experimental control over the task environment.

Here, we compare the performance of two deep reinforcement learning (RL) agents and a rhesus macaque in a pursuit avoidance task that is complex enough to challenge to state-of-the art reinforcement learning models but simple enough to train non-human primates to high levels of performance. In this paradigm, the agent (used henceforth to refer to both the non-human primates and deep RL agents) is faced with continuous decisions about where to move based on the position of a predator that the agent must avoid and the position of a prey that the agent must catch to be rewarded. The prey and predator follow cost-driven movement policies that involve repulsion from and attraction to the agent, respectively. These highly interactive dynamics, paired with sparse rewards and punishments contingent on avoiding the predator and catching the prey, present a significant challenge to reinforcement learning models. Here we train a deep Q-learning network, as well as a deep actor-critic model to perform this task, and compare the two artificial agents against the non-human primates with respect to (a) overall performance on task, (b) learning dynamics, (c) behavioral strategies as a function of prey and predator position, and (d) the complexity of the internal model of the task that is reflected in the agent’s decision-making behavior. We discuss implications of this analysis for the development of future artificial agents, as well as for understanding the latent cognitive variables underlying decision-making processes in non-human primates.

## 2 Methods

### 2.1 Virtual pursuit-avoidance task design

We designed the virtual pursuit-avoidance task based on an existing visual experimental design (*5*). In this design, the agent controls the position of an avatar (yellow circle in Fig. 1). The game determines the predator and prey’s next positions at each step according to attraction and repulsion force functions, along with a cost contour map over the pixel field (1920 by 1080 pixels). The field cost is higher along the perimeter to mitigate cornering tactics. The color of the predator and prey objects encodes their maximum speed. The agent is always yellow. Unless otherwise stated, the starting points of the objects and their colors are randomized trial-by-trial, with a minimum starting distance to the agent of 400 pixels.

**Figure 1:**
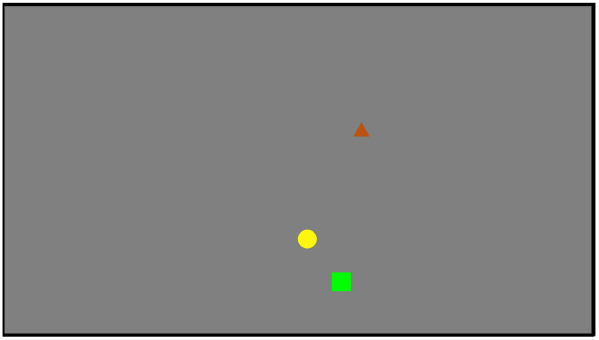
The visual environment. The agent (yellow circle), predator (red triangle), and prey (green square) are shown.

The macaque interacted with the environment through a joystick providing proportional inputs in two dimensions (−1 ≤ *x,y* ≤ 1). The deep RL agents interacted through an environment API. The API provides an (`observation, reward, done`) tuple each trial step and accepts an action vector. The observation consisted of the color values and x-y coordinates of the agent, predator, and prey. Steps were assigned a small penalty, predators a large penalty, preys a large reward, and timeouts (max steps reached) a medium penalty. The action vector was agent-dependent (see below). The original environment was implemented in Matlab using Psychtoolbox (*6*) as an interactive simulation with joystick-driven input. We implemented a new environment in Python using PsychoPy (*7*) for training the RL agents. The agent interface conforms to the de facto standard defined by the OpenAI Gym project (*8*).

### 2.2 Macaque training and evaluation

A male adult Macaca mulatta (8 year old) was trained using a multi-stage curriculum. The goal was to learn the following in sequence: 1) touching the joystick produces reward; 2) moving the joystick controls the position of a circle (agent’s avatar) on the screen; 3) overlap of the circle and a square (prey) on the screen provides a reward; 4) objects move; 5) color indicates the amount of reward as well as maximum speed of the prey; 6) a triangle (predator) will pursue the circle and overlap results in a penalty.

### 2.3 Deep reinforcement learning agents

The deep Q network implemented Q-learning, a form of off-policy temporal-difference learning that estimates an action-value function through iterative updates. This function estimates the expected value of all future rewards for a given state and action (Eqn. 1).

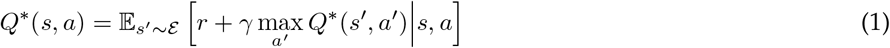

Given the size of the state space, this function was approximated using a fully-connected deep neural network (*1*). The model performed online RL using two copies of the same network - the policy network and the target network. In our “double DQN” (*9*), the policy network updated the action-value function as it replayed prioritized experiences from memory. Periodically its parameters were loaded into the target network, which took action in the environment. Replay was prioritized by the magnitude of an experience’s TD error (*10*). Epsilon greedy exploration was used in the bootstrapping process. A short sequence of observations were “stacked” to provide the input state to the network. The network output was a 1-hot state vector of 9 possible actions (cardinal and sub-cardinal directions and none). We used our own Python implementation.

Soft Actor-Critic (SAC) (*11*) is an off-policy actor-critic agent that simultaneously maximizes expected return and policy entropy:

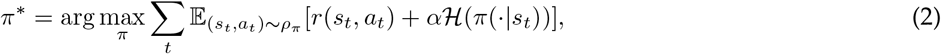

SAC automatically uses approximating dual gradient descent to tune the parameter *α* that determines the importance of entropy. The critic part of SAC uses the minimum of two soft Q function approximators to counter positive bias when computing gradients. It also uses a target soft Q function and a replay pool, like DQN. SAC outputs continuous [−1, 1] actions, emulating joystick input. We used the Rlkit implementation of the agent (*12*).

### 2.4 Deep RL agent training and evaluation

The deep RL agents were trained on randomly-initialized episodes. The agents executed until one of the termination conditions - catch, caught, or maximum steps - was encountered. The agents were trained using backpropagation until catch performance stopped improving, somewhere between 6,000 and 10,000 episodes. DQN used epsilon greedy exploration, starting at 1.0 and ending at 0.25; epsilon was 0 for evaluation. SAC, which targets maximum entropy for its policy, does not inject explicit exploration noise; instead, it uses the mean action.

DQN used a frame skipping technique, processing every 3rd frame, to lower the training time without eliminating useful motion information. Memory replay capacity was 100,000 episodes, which were replayed at 128 episodes per step based on rank-based importance sampling.

SAC processed every frame and its replay memory capacity was 1,000,000 episodes, not prioritized, with a batch size of 1024. Notably, SAC achieved high performance on the task using default hyperparameters. In contrast, DQN was extremely sensitive to many of the hyperparameters listed above.

The agents were evaluated on a different set of randomly initialized episodes (> 10,000). The environmental parameters used (screen size, object size, object speed, etc.) matched the parameters used with the macaque. The network model parameters were frozen during this phase (i.e., no backpropagation).

## 3 Results and Discussion

### Qualitative comparison of training and learning characteristics

The macaque was trained on the task for 70 days and 10,850 trials, after finishing the previous curriculum stages. The agents reached peak performance in a similar number of trials. Training took 2-10 hours on single-CPU machines. The training curves are shown in Fig. 2.

**Figure 2:**
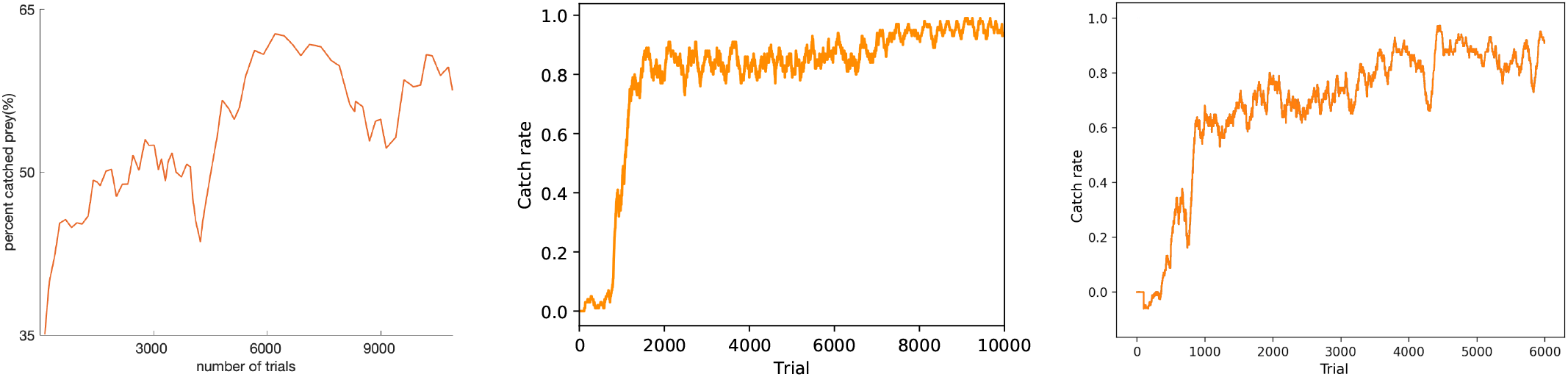
Training performance curves for the macaque (left), SAC agent (middle), and DQN (right). The macaque performed an average of 155 trials per day (10,850 total). The catch rate performance for the agents is averaged over a trailing sliding window of 100 episodes.

### Overall performance on the task

We compared overall performance after the macaque and the agents had achieved peak performance on the task. The results are shown in Table 1. Both agents greatly outperformed the macaque on the task. The superior performance of the RL agents is not attributed to a lack of attention or motivation by the macaque, a fact supported by the macaque’s high performance on other types of trials (e.g., one prey, no predator) (*5*).

**Table 1:** Peak performance on the task for the macaque (MK), the DQN agent, and the SAC agent. Termination rates and the average number of steps to termination are shown.

| Player | Catch |  | Caught |  | Timeout |  |
| --- | --- | --- | --- | --- | --- | --- |
|  | Percentage | Mean steps | Percentage | Mean steps | Percentage | Mean steps |
| MK | 66.67 | 168 | 33.33 | 169 | 0.26 | 1200 |
| DQN | 98.5 | 154.8 | 1.29 | 88.5 | 0.21 | 1200 |
| SAC | 99.99 | 48.65 | 0.01 | 43.03 | 0 | N/A |

### Evidence that the macaque and the RL agents model the physics of motion

We analyzed the step-by-step trajectories for evidence that the agents had learned to instantaneously predict prey and predator movement by modeling 1st, 2nd, and/or 3rd order equations of motion. Previous work (*5*) had found evidence that macaques learn latent representations of these motion mechanics. The generative model described in (*5*) and depicted in Fig. 3 was used. In summary, the macaque’s behavior was best explained by 2nd order physics, qualitatively matching the findings of (*5*) even though the current study adds a predator to the environment. In contrast, the behavior of both artificial agents was better explained by the 1st order prediction model. One explanation for this simplified modeling is that the agents are unencumbered by mechanical inertia, in contrast to the macaque, and execute more precisely-aimed actions (on average; see below).

**Figure 3:**
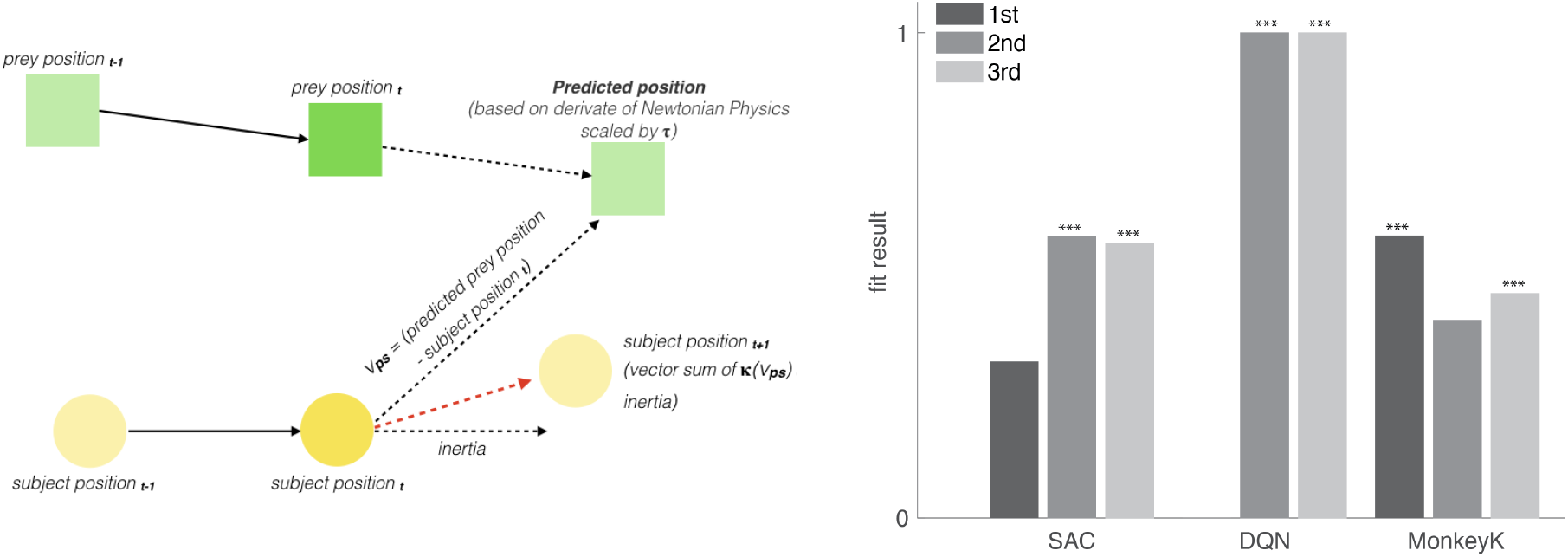
Fitting generative models of motion physics. The agent (yellow) moves based on past prey movement information (position, velocity, acceleration). The pursuit vector is scaled by a force parameter *κ*; the resulting position of the subject is then computed by summing the pursuit vector and the joystick movement inertia. Once the target location is set, an action is taken. Performance of each motion model was compared trial-by-trial and averaged across trials. Lower values indicate better fits, with significance values indicated for within-agent comparisons (stars indicate *p* < 0.001).

### Evidence that the macaque and the RL agents take similar/different actions depending on context

We analyzed the actions taken by the macaque and agents under the same environmental conditions (i.e., the same avatar, predator, and prey locations, and the same values). To do this, we presented the agents with an observation that the macaque had seen and taken action on. We then measured the difference in angle of action input by the macaque and agent. We binned each measurement based on the distance of the avatar to either the predator or the prey. The results are shown in Fig. 4.

**Figure 4:**
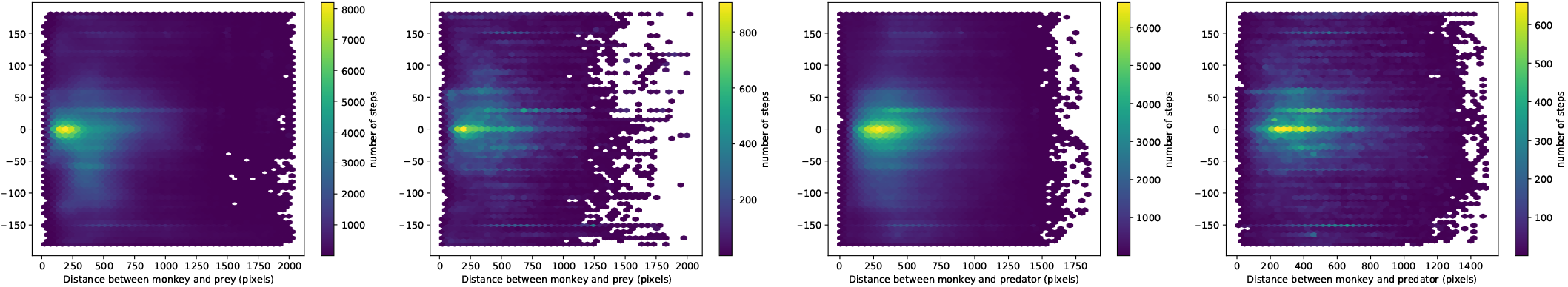
Action angle agreement as a function of distance to prey and predator. The color indicates the number of episode steps per bin. From the left: SAC for prey distance, DQN for prey distance, SAC for predator distance, and DQN for predator distance.

For SAC, there is strong angular agreement when the avatar is within a couple hundred pixels of the prey (prey diameter is 60). The DQN exhibits its best agreement under this circumstance as well, but the agreement is less pronounced and coherent. This may be due to observed action oscillations due to the DQN’s limited one-hot action space (versus the SAC’s continuous action space). A similar pattern of agreement is seen as a function of predator proximity, but the pattern is more spread out because the typical agent-predator distance spans a greater range.

### Trajectory analysis

We analyzed whether the agents were more effective than the macaque when initialized from steps along trajectories in episodes that the macaque had played to termination. We conditioned the analysis on outcome: 50 episodes that terminated in prey capture and 50 that terminated in being caught by the predator. At every step, we initialized the environment and had the agents play to assess relative performance in terms of outcome and steps to termination. The results are shown in Fig. 5 (agent outcomes not shown). In general, the agents capture the prey in significantly fewer steps, but with less advantage when initialized deeper into the trajectory (closer to the prey/predator). The DQN in particular performs worse when initialized close to the prey, possibly due to action space oscillations and an inability to immediately compensate for them, given the proximity. Furthermore, agents were caught much less frequently than the macaque, even when initialized with only 10% of the trajectory remaining (i.e., in close proximity). In these cases, the number of remaining steps is higher, primarily due to them escaping and eventually pursuing the prey. Both agents survived much longer than the macaque, and often escaped to later capture the prey, when initialized close to the predator.

**Figure 5:**
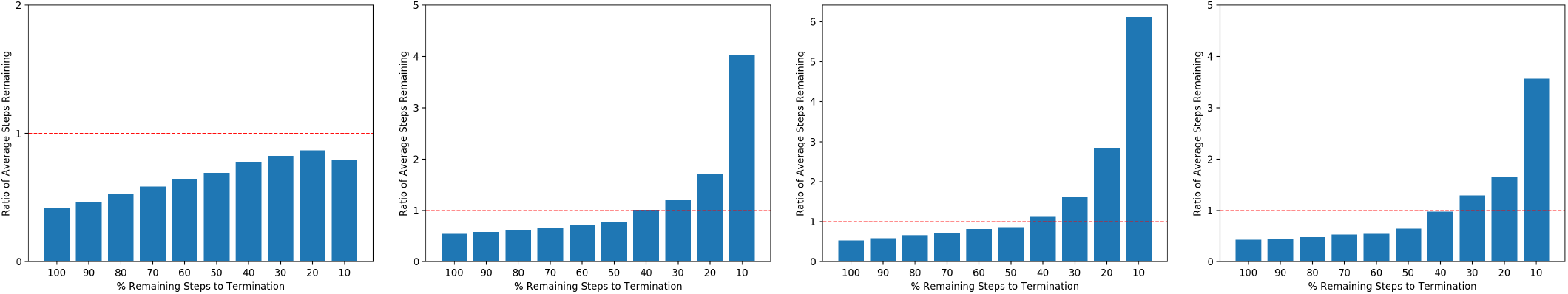
Trajectory analysis. From the left: SAC initialized on macaque trials that ended in prey capture; DQN for prey capture trials; SAC initialized on macaque trials that ended in being caught by predator; DQN for caught by predator trials. The RL agents were initialized progressively at steps within the trials and run to termination. Lower is better for trials that end in prey capture by agent (i.e., nearly all trials, not shown). Red line indicates that the macaque and agent would terminate in the same number of remaining steps.

## 4 Conclusion and Future Work

We observed that the RL agents learned as fast and to a higher level of proficiency than macaques (*5*), in contrast to reports of similar agents learning to play games much more slowly than humans. This may be due to the simplicity of our particular environment, which limits complexity to the mechanics of motion and relatively simple interactions. Peak agent performance was near perfect and they were much more efficient at achieving successful outcomes. In general, the agents exhibited a tendency to take the same action as the macaque when put into identical circumstances. However, over a series of steps the outcomes played out quite differently. The reason for this remains inconclusive. Motor control inertia and less motor precision by the macaque may explain this.

A growing number of studies suggest that humans (*13*), non-human primates (*5, 14, 15*), as well as other mammals (*16,17*) develop predictive models of the environment to guide their decision making. In line with these results, we observed that behavior of the non-human primate can be well explained by a predictive model based on 2nd order Newtonian physics. In contrast, the behavior of trained artificial agents was best explained by a simpler model of 1st order Newtonian physics. The latter confirms recent criticisms of AI, arguing that the learning of naturalistic agents is guided by a domain-general knowledge of intuitive physics, whereas artificial agents often rely on the acquisition of domain-specific knowledge to solve the particular task with which they are confronted (*4*), without necessarily learning more general characteristics of the environment or strategies that may exploit those for other purposes. The lack of such inductive biases can result in higher performance on one task at the expense of learning efficiency and generalization performance (*18*).

The experimental setup in this work provides opportunities for a more systematic comparison of artificial and natural agents, including the opportunity to measure neural correlates of performance. The latter can be used to test for latent variables predicted by the computational models. The computational models, in turn, can be used to test the effects of constraints on processing that may help explain performance of the natural agents. For example, one candidate explanation for the lower performance of the non-human primate is a limitation in the ability to accurately perceive and/or process information about the predator and prey at the same time, thus requiring cognitive control to determine where to allocate attention (and even fixation) at a given time. Recent work suggests that such limitations take the form of a cost that attaches to the allocation of cognitive control (*19, 20*). The paradigm and computational models described here provide a platform for testing hypotheses about the nature of such costs, and the mechanisms used to evaluate them and allocate control.

